# The workings and failings of clustering T-cell receptor beta-chain sequences without a known epitope preference

**DOI:** 10.1101/318360

**Authors:** Pieter Meysman, Nicolas De Neuter, Sofie Gielis, Danh Bui Thi, Benson Ogunjimi, Kris Laukens

**Affiliations:** Antwerp Unit for Data Analysis and Computation in Immunology and Sequencing (AUDACIS), University of Antwerp, Antwerp, Belgium; ADREM data lab, University of Antwerp, Antwerp, Belgium; biomedical informatics research network Antwerp (biomina), University of Antwerp, Antwerp, Belgium; Antwerp Center for Translational Immunology and Virology (ACTIV), Vaccine & Infectious Disease Institute (VAXINFECTIO), University of Antwerp, Wilrijk, Belgium; Centre for Health Economics Research & Modeling Infectious Diseases (CHERMID), Vaccine & Infectious Disease Institute (VAXINFECTIO), University of Antwerp, Wilrijk, Belgium; Department of Pediatrics, Antwerp University Hospital, Edegem, Belgium

## Abstract

The T-cell receptor is responsible for recognizing potentially harmful epitopes presented on cell surfaces. The binding rules that govern this recognition between receptor and epitope is currently an unsolved problem, yet one of great interest. Several methods have been proposed recently to perform supervised classification of T-cell receptor sequences, but this requires known examples of T-cell sequences for a given epitope. Here we study the viability of various methods to perform unsupervised clustering of distinct T-cell receptor sequences and how these clusters relate to their target epitope. The goal is to provide an overview of the performance of various distance metrics on two large independent T-cell receptor sequence data sets. Our results confirm the presence of structural distinct T-cell groups that target identical epitopes. In addition, we put forward several recommendations to perform T-cell receptor sequence clustering.

## Background

T-cells constitute an important part of the adaptive immune system against invasive pathogens and pathological cells. These T-cells are capable of identifying self from non-self antigens and triggering the adaptive immune response. At a molecular level, T-cells carry a protein complex called a T-cell receptor (TCR) on their cell surface, which is able to bind antigen epitopes presented by the major histocompatibility complex (MHC) molecules on host cells. Two types of MHC molecules exist, namely class I MHC and class II MHC molecules. Class I MHC molecules typically bind shorter epitope peptides that originate from within the cell. The Class I MHC molecules are recognized by CD8+ T-cells, the so-called cytotoxic T-cells. Broadly these cytotoxic T-cells will target cells that are presenting non-self peptides which may be indicative of a viral infection within the cell or a tumor cell. Class II MHC molecules bind longer epitopes that are typically considered to be extracellular in origin, which are bound by CD4+ T-cells. The TCR complex on these CD4+ or CD8+ T-cells itself is composed of two protein chains, an alpha chain and a beta chain. Each chain is the result of a recombination event during the maturation of the T-cell in the thymus (Bassing *et al*., 2002). This features a somatic rearrangement of noncontiguous variable (V), diversity (D), and joining (J) region gene segments for the beta chain, and V and J segments for the alpha chain, supplemented by the random addition or removal of nucleotides at the joints. This creates a highly variable receptor that is unique to each T-cell clone. The region of highest variability is the third complementarity-determining region (CDR3) region, which is also the primary region for binding, and therefore recognizing, epitope peptides. A large number of potential combinations is possible, allowing the body to create T-cells that recognize a large number of different epitopes, and in turn a large number of pathogens and malignant cells (Robins *et al*., 2010).

Recent technical improvements within high-throughput sequencing technologies have created several experimental methods to perform so-called TCR sequencing (Robins *et al*., 2009). Through this technique, fixed primers can be used to selectively sequence the variable CDR3 region within the TCR RNA or DNA for an entire sample of T-cells. The fact that the primers are specific for T-cells with recombined TCR sequences means that purification from whole blood is not strictly necessary. TCR sequencing now enables us to acquire an overview of an individual’s entire repertoire, providing an overview of the millions of different TCR sequences composing this part of the adaptive immune system. Several tools exist to process raw TCR sequencing data into a quantitative list of TCR sequences with their likely VDJ recombination events typed (Alamyar *et al*., 2012; Gerritsen *et al*., 2016; Bolotin *et al*., 2015; Thomas *et al*., 2013). It has already been shown that TCR sequence data is sufficient to predict cytomegalovirus seropositivity with high accuracy (Emerson *et al*., 2017; De Neuter, Bartholomeus, *et al*., 2018; Pogorelyy *et al*., 2018). There is thus a vested interest within the scientific community to further develop these techniques and analyse methods to process TCR data (Miho *et al*., 2018).

One key remaining question concerns how the epitope recognition of the TCR complex works on a molecular level. In the past year, three methods have been published that created a computational model to predict binding between an epitope and a TCR sequence based on a set of known interactions (De Neuter, Bittremieux, *et al*., 2018; Dash *et al*., 2017; Glanville *et al*., 2017). All three approaches work in a supervised fashion. A training data set with TCR sequences known to bind a specific epitope is supplied and the model attempts to predict which other TCR sequences may bind the same epitope. Each of these methods independently reported high performance, but as they were all published within the same time window, they were not compared. Moreover, each of these approaches relies on known TCR sequences for a given epitope. Despite ongoing curation and collection efforts (Shugay *et al*., 2017), high quality epitope-specific TCR sequences remain rare, as they require costly experiments often involving dedicated MHC tetramers for each epitope under investigation. Thus, there is only TCR data for a few hundred epitopes, while the real epitope space potentially contains every peptide sequence from length seven until thirty. Furthermore, the majority of TCR data that is currently available are repertoire-wide screens at an individual level. These are tens of thousands of TCR sequences without any knowledge of the specific epitope that they target, and only some information regarding the individual of origin. There is thus a clear need for an unsupervised approach that can group TCR sequences binding the same, unknown, epitope. In addition, the majority of TCR sequencing studies currently being published are focused on the TCR beta chain. It is a non-trivial problem to sequence both the alpha chain and the beta chain of the TCR in such a manner that they can be assigned together as originating from a single T-cell clone. Several solutions exist, including single T-cell receptor sequencing, but they remain far from commonplace (Redmond *et al*., 2016; Stubbington *et al*., 2016). Finally, many fundamental questions remain regarding the binding between a TCR and its epitope. Reason and research indicates that similar TCR sequences should recognize similar or the same epitopes. However, how dissimilar TCR sequences can be before this is no longer the case is not known. Moreover, there is no consensus on how to define the similarity between two short TCR CDR3 amino acid sequences.

In this paper, we compare several unsupervised approaches to cluster TCR sequences based on their similarity. In particular, unsupervised versions of the supervised techniques that have proven successful in recent publications are included. The idea is that the features used by these methods should have captured important properties of the TCR if they have high performance in the supervised approach. In this manner, we evaluate the clustering of epitope-specific TCR sequences based only on the beta chain sequence features.

## Methods

### Data

Only human TCR data was considered for these analyses. While there is also a wealth of mice TCR data, distinguishing TCRs from different species was not a goal of this study. We used two datasets: a smaller dataset from a single study where all TCRs originate from a limited set of individuals and a larger dataset from the curated VDJdb, which collects TCR data from many different studies and therefore the TCR sequences originate from many different individuals (Shugay *et al*., 2017). In both cases, only the V-region, J-region and CDR3 amino acid sequence was used of each TCR beta sequence.

The smaller dataset contains TCR sequences targeting one of three epitopes originating from an infectious disease, namely the influenza M1 epitope, the Epstein-Barr virus BMLF epitope and the CMV pp65 epitope. We term this dataset the “Dash dataset” and it contains a total of 412 unique TCR beta-chain sequences. Note that this is also the dataset for which the supervised version of the GapAlign method was designed (Dash *et al*., 2017).

The larger dataset was downloaded from the VDJ database on the 14^th^ of September 2017. Only TCR sequence - epitope relationships were considered with a vdj.score higher than 0, thus removing those deemed by the VDJ database as unreliable. This dataset contained 2065 TCR beta sequences with 100 unique epitopes presented on 20 unique MHC molecules (17 MHC-I and 3 MHC-II).

### Distance measures

#### 1. Length-based distance

The distance between two TCR sequences is defined as the difference in number of amino acid in the CDR3 region between them, where the CDR3 region is defined as the amino acids flanked by and including the 104C and 118F residues, as per IMGT notation (Lefranc and Lefranc, 2002). This distance measure is solely used as a comparative baseline. As we will show, the TCR length is a confounding factor within many distance measures. In some cases, grouping TCR sequences based on length may already provide viable clusters targeting specific epitopes.

#### 2. GapAlign score

This distance measure is derived from the supervised approach used in (Dash *et al*., 2017). It is based on a sequence alignment problem where a single gap is allowed. The sequence scores are derived from a flattened BLOSUM90 matrix. The original use was within a k-nearest neighbour approach and thus the same scoring scheme can be readily adapted for unsupervised use. To allow comparison, the original code published by Dash et al. was used to generate the scores. In this case only the beta-chain CDR3 scoring scheme was used, instead of the combination of both the alpha and beta chains. The default scoring scheme was applied, thus gap penalties are set to 8. The code was updated from python 2.7 to python 3 for integrative purposes.

#### 3. Profile score

This distance measure is based on the physicochemical differences between two TCR sequences and is derived from the approach taken by (De Neuter, Bittremieux, *et al*., 2018). The basicity, helicity and hydrophobicity values for each amino acid are Z-normalized and used to construct a profile of the full TCR beta CDR3 sequence. The longest TCR sequence is truncated along either side to match the shortest sequence. The distance between the two profiles is then calculated by the weighted Euclidean distance, with a higher weight for the central positions and decreasing in a linear fashion to the edges of the CDR3 profile. The final score is then the sum of the distances for each of the three physicochemical profiles.

#### 4. Trimer scor

This distance measure calculates the percentage of similar amino acid trimers between two TCR sequences and is derived from the approach by (Glanville *et al*., 2017). In addition, this is similar to other trimer-based methods, such as (Thomas *et al*., 2014). As TCRb CDR3 sequences often start with CAS and end with YFF, some trimers are more informative then others. Therefore, a weighting scheme was introduced. This is also in line with the original supervised approach where trimers were used which were statistically overrepresented in a TCR data set targeting a specific epitope or antigen, versus one that did not (Glanville *et al*., 2017). The weighting scheme was based on the combined CD8+ TCR repertoires of all the data collected within the study of (Ogunjimi *et al*., 2017). These were full repertoires without any prefiltering on epitope preferences and can therefore be considered as representing the modal TCR diversity. The occurrence fraction of each trimer was calculated. Each trimer is then assigned a score of 1 subtracted by this fraction. Thus, an unseen trimer is assigned a value of 1, while the common CAS trimer only receives a score of 0.17. The score of each shared trimer is summed between the two TCR sequences. As distance measures have to be lower for more similar samples, this score was normalized for the number of trimers in the shortest sequence and this sum was subtract from 1. In this manner if two TCR sequences share rare trimers the distance score will be smaller than if they share more common variants.

#### 5. Dimer score

This score is equivalent to the trimer score but uses amino acid dimers instead of trimers.

#### 6. Levenshtein distance score

This score is defined as the Levenshtein distance (also known as the edit distance) between the TCRb CDR3 amino acid sequences. The score corresponds to the minimum number of mutations, deletions and insertions needed to transform the first sequence into the second sequence. This distance has been used on several occasions for clustering TCR sequences (Tickotsky *et al*., 2017; Madi *et al*., 2017).

#### 7. VJ edit distance

This score represents the similarity of the V- and J-region used to assemble the TCR beta-chain. This score is the Levenshtein distance between the amino acid content of the original V-region sequence and original J-region sequence summed. This is the only distance measure within this study that is not only based on the TCR CDR3 amino acid sequence and can be considered as complementary to each of the other approaches.

### Clustering algorithm

The DBSCAN algorithm from the python scikit-learn library (0.19.0) was used to cluster the TCR sequences based on the generated distances measures. The minimum amount of samples per cluster was set to two. The advantage of the DBSCAN algorithm is that it does not require one to specify the number of clusters in advance. Thus, we do not need to introduce any prior knowledge on the number of clusters that may be presented in the dataset. This is comparable to the real-life case where we are given a dataset of TCR sequences without any known epitopes. DBSCAN will group samples based on a fixed distance, defined in advance. This allows us to try different similarity thresholds on the TCR sequences and evaluate their use. In addition, this makes the algorithm particularly robust against outliers, which can be expected to be very common in TCR data. The used threshold is based on a fixed point within the distribution of each distance measure. It is allowed to vary between 0.1% of the lowest reported distances to 40% of the lowest reported distance to establish a wide range of potential clustering solutions.

Note that the length-based distance can only work with the DBSCAN algorithm on a single threshold, namely a value between 0 and 1. Higher than this value the DBSCAN algorithm will create a single cluster as there is a TCR sequence of every length between 10 and 20 contained in both datasets.

### Clustering performance measures

Evaluating the quality of an unsupervised clustering approach is a non-trivial problem, even if the ground truth is known. Especially for the DBSCAN approach, which allows samples to be left unassigned to any cluster and have clusters of different sizes. We define the following commonly used supervised performance metrics for use within this setting, illustrated in figure 1:

**Figure 1:**
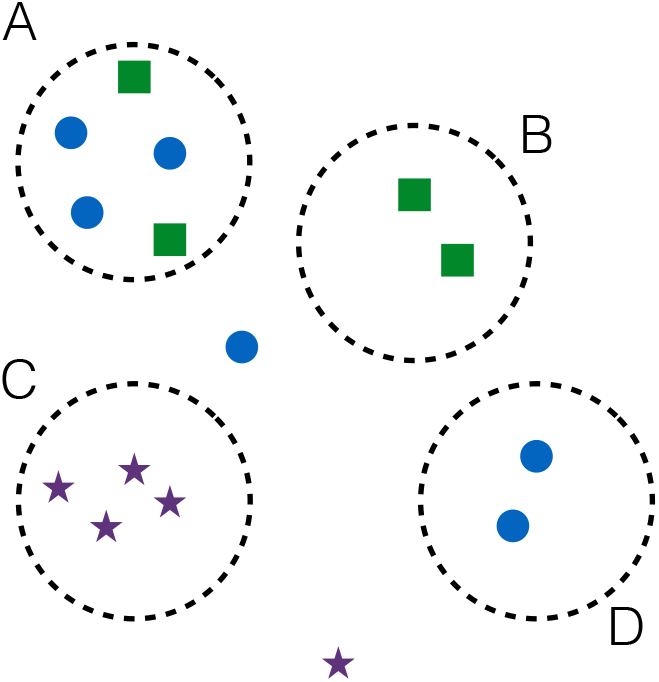
Illustrated example of possible grouping results for 15 TCR sequences known to bind three different epitopes (circles, squares and stars) in four different clusters (A, B, C and D). The different metrics will then have the following outcomes. Recall: the percentage of TCR sequences assigned to a cluster. Only two are not in a cluster, thus the recall will be equal to 13/15. Accuracy: each epitope is assigned a true cluster. In this case, cluster A for the circles, cluster B for the squares and cluster C for the stars. There are equal amounts of squares in cluster A and B, but preference is given to cluster B for the squares as cluster A has more circles. Cluster D is considered as a false positive cluster as cluster A has the most circles and is therefore assumed to be the true cluster where all circles should be grouped. The accuracy is then (3 + 2 + 4) / 15, based respectively on the number of correct assignments in clusters A (3), B (2) and C (4). Precision: for each cluster, the most common epitope is considered a true positive irrespective of its co-occurrence in other clusters. This sum is then divided by the amount of TCR sequences that are assigned to a cluster. The precision will then be 3 + 2 + 4 + 2 / 13, based on the circles in cluster A (3), the squares in B (2), the stars in C (4) and the circles in D (2).

#### 1. Accuracy

In this instance, accuracy is defined as the fraction of TCR sequences recognizing the same epitope that are assigned to a single cluster. If two or more clusters contain TCR sequences assigned to that epitope, only the cluster containing the most is counted as being the true cluster for that epitope. If two epitopes end up with the same true cluster, those with the most TCR sequence is given preference and the other epitope is assigned to the next cluster in line. In this manner there is only a single cluster assigned to each epitope. This measure is therefore equivalent to the supervised accuracy as it determines how many TCR sequences are assigned to the true cluster determined for their epitope. This measure provides an indication for how well a method can place all TCR sequences targeting the same epitope in a single cluster.

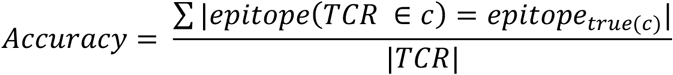

#### 2. Recall

In this instance, recall is defined as the fraction of TCR sequences that have been assigned to a cluster. This measure gives us an indication on the percentage of TCR sequences that can be grouped together given a certain clustering threshold.

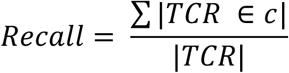

#### 3. Precision

In this instance, precision is defined as the fraction of TCR sequences within a single cluster to be targeting the same epitope. The most common epitope is considered to determine the fraction. In this case, multiple clusters can be evaluated on the same epitope. This measure therefore provides an indication of the purity of each cluster, i.e. if TCR sequences are grouped together how likely they are to all recognize the same epitope.

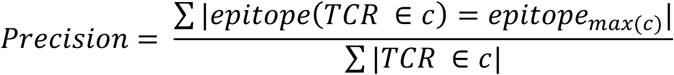

However, as the used datasets are unevenly balanced with respect to the number of TCR sequences for each epitope, it is important to establish a comparative baseline. For example, imagine a dataset where 90% of the TCR sequences bind the same epitope. Any random configuration of clusters will have an average of 90% sequences binding the same epitope. Any method will report high accuracy and precision on this dataset even if it is not better than random. Therefore, after every clustering step, we randomize the group assignments fifty times to create a baseline for comparison. Note that this may differ from method to method as some methods result in many smaller clusters while others have fewer larger clusters.

All code, both for the generation of the distance measures and the calculation of the performance metrics, is available on github: https://github.com/pmeysman/TCRclusteringPaper

The scripts are designed so that they can be readily extended with new distance measures, which can then be applied to the same datasets and compared to the ones described in this study.

## Results and discussion

### CDR3 length is a confounding factor in many TCR sequence distance measures

We first applied the six distance measures to the Dash dataset containing 412 TCR sequences. A distance score is assigned to each possible pair of TCR sequences, which results in a distance matrix of 412 × 412 for each measure. Prior to applying the clustering algorithm to these distance matrices, we investigated the relationships that exist within the distance scores themselves. All distance scores were concatenated into a single vector and compared with a principle component analysis, whose results can be found in figure 2A. As expected the VJ edit distance is a clear outlier and is independent from the other measures as it is based mostly on a different part of the TCR sequence. The loadings assigned to all other measures, including the TCR sequence length, cluster closely together. This indicates that a large distance in one score corresponds to a large distance for another as well. The analysis was therefore rerun without the VJedit measure, resulting in figure 2B. In this instance, there is a clear grouping between the Dimer and Trimer distance measure, which can be expected as common trimers are intrinsically made up of two common dimers. The GapAlign, Levenshtein and Profile methods are also suggested to be related. These methods can be considered as variants of sequence alignment on the full CDR3 sequence and therefore likely provide similar scores. The Length measure is less grouped with the others compared to the case when the VJedit measure was included. This suggests that there is information captured by the various distance measures that is independent from the CDR3 Length difference.

**Figure 2:**
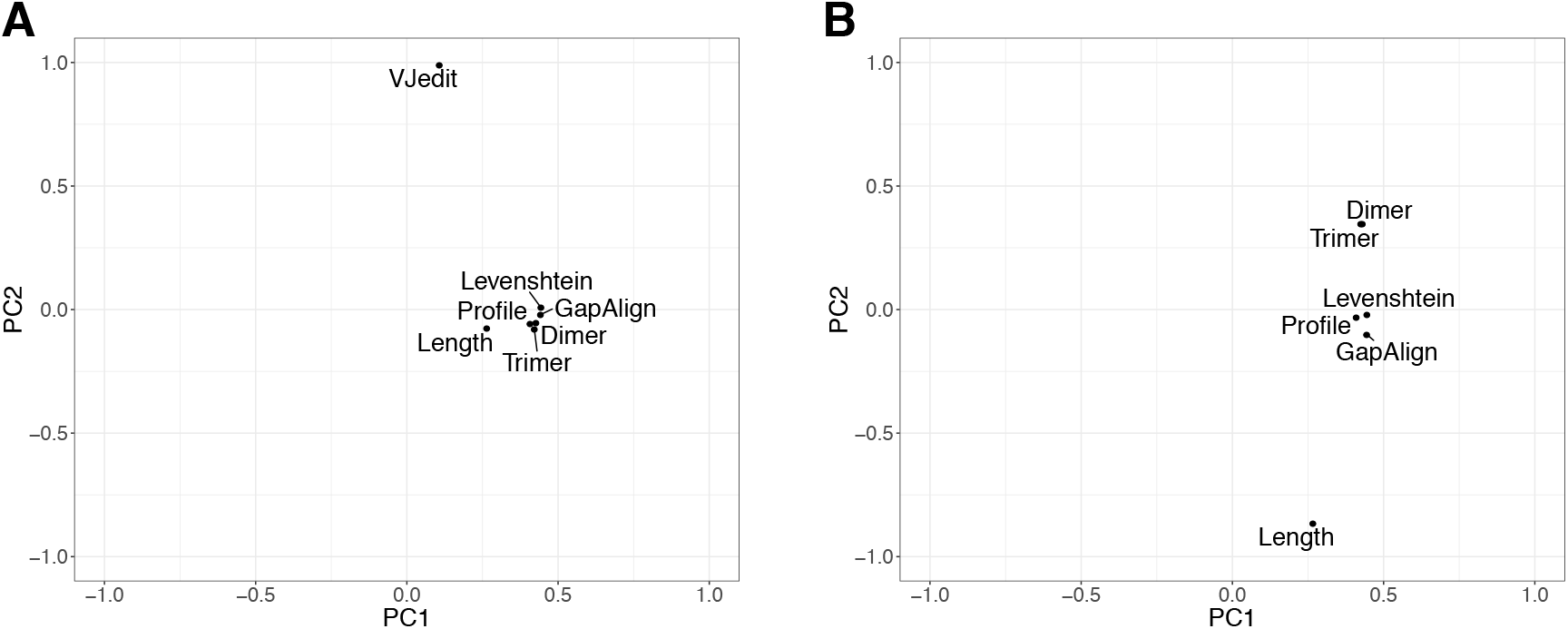
PCA loading plots for the concatenated distances matrices for the different distance measures. A. With the VJedit distance. B. Without the VJedit distance.

The VDJdb dataset contains 2065 TCR sequences and with this larger set, we can explore if there are any natural groupings within the TCR beta chain sequences that are highlighted by the different distance measures. The results from PCAs applied on the distance matrix of each measure can be found in figure 3. The length distance measure exists as a gradient from longer to shorter CDR3 sequences. As made clear from the PCA score plots, the length is unable to make a distinction between CDR3 sequences of the same size. The GapAlign distance measure presents as a dense cloud. However, the first principle component matches strongly with the CDR3 amino acid length. This is the result of the introduced gap penalty, as a sequence that is longer by one amino acid will be punished worse than a sequence that has the worst possible single amino acid substitution. This is a common aspect of any alignment approach, and removal of the gap penalty is not a viable option due to the differences in TCR CDR3 length. The Profile distance measure is also influenced by the sequence length, but as suggested by the PCA score plot, to a lesser degree. This is likely as each CDR3 sequence is centred on the central amino acid and additional amino acids are not considered of the sequences differ in length. However, there are two clear groupings within the Profile PCA score plot. Closer investigation reveals that these are the CDR3 sequence with an even and an uneven length. As the central position is shared for even sequences but not for the uneven sequences, the alignment and the scoring has to work in a different fashion. This results in a split along the second principle component. The Trimer and Dimer distance measures feature similar PCA score plots. Each of the two first principle components has an independent high scoring group and the remainder has a score of around zero. Each high scoring group is typed by a common pentamer or hexamer at the end. For example, high scores are assigned in PC1 of the trimer distances to short CDR3 sequences that end in SYEQYF. This corresponds to the start of the TRBJ2-7∗01 sequence. These are thus all sequences that are derived from this specific J gene, without removal of any nucleotides during the recombination event. In the same way high scores in PC2 are assigned to CDR3 sequences ending in YNEQFF, corresponding to the start of the TRBJ2-1∗01 gene. The Levenshtein distance has similar trends to the aforementioned measures. There is a clear trend along PC1 related to the length of the CDR3 sequence. In addition, PC2 seems to divide the TCR sequences into three groups. These three groups correspond to three common J-genes, namely TRBJ2-1∗01, TRBJ2-2∗01, TRBJ2-3∗01 from top to bottom. The VJedit distance features a large number of smaller groupings, corresponding to common V or J gene combinations.

**Figure 3:**
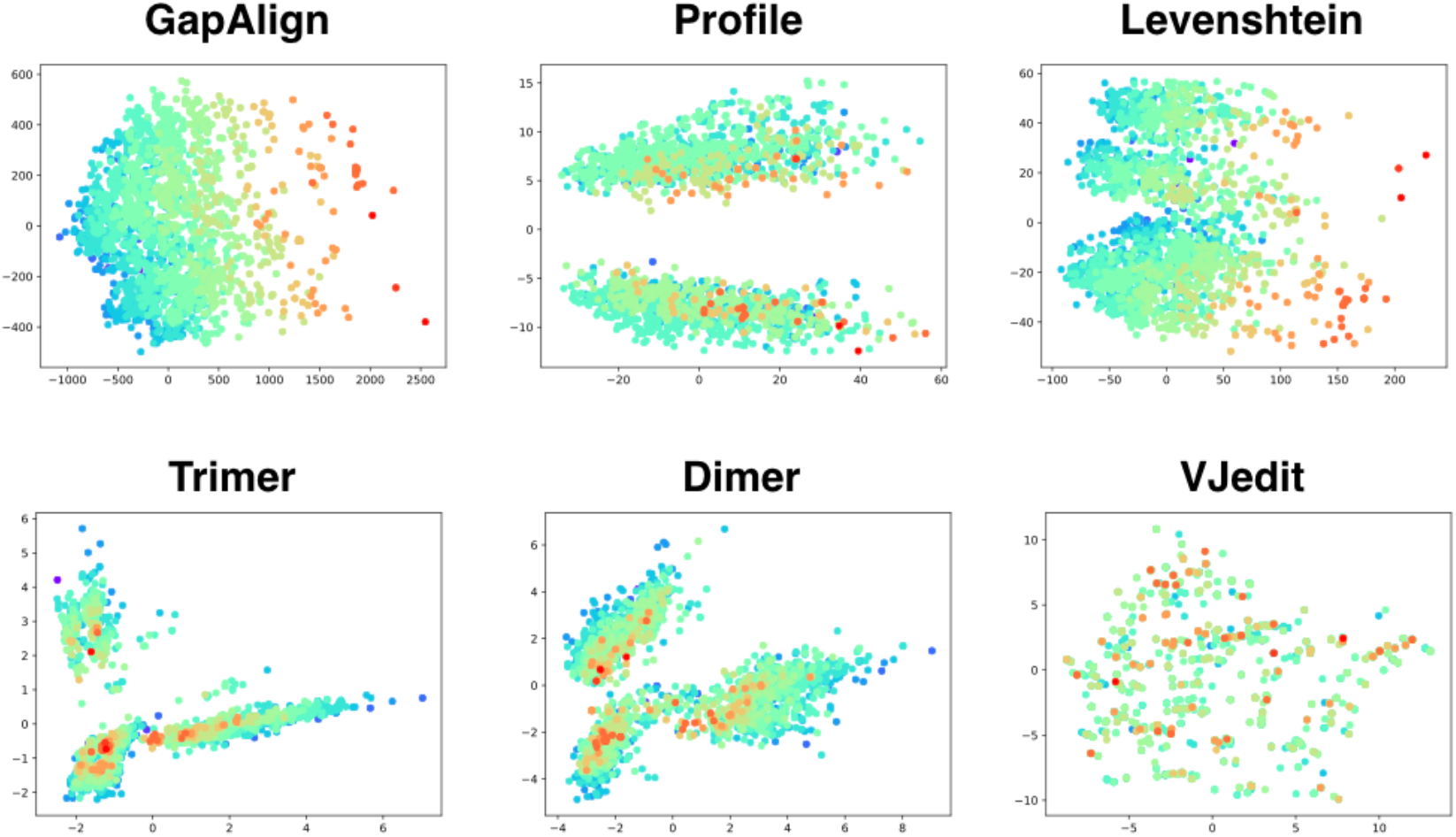
PCA score plots for each distance measure, with the first principle component (PCI) as the horizontal axis and the second principle component (PC2) as the second axis. Each point is a single TCR sequences from the VDJdb dataset. Colors represent the length of the TCR, with red as the longest and purple as the shortest.

### Most methods establish a high precision but struggle with high accuracy

While we have uncovered several relationships that exist within each distance measure, none provide any indication on which distance measure performs best at grouping TCR sequences that target the same epitope. Indeed, scores that indicate strong similarity between TCR sequences that have the same length or are derived of the same J gene, may provide strong performance as they are still grouping highly similar TCR sequences together.

As a first test, a ROC plot can be used to visualize the likelihood that two TCR sequences that bind the same epitope will receive a higher score than two TCR sequences that do not. This is possible in this unsupervised context since we know the ground truth for each of the TCR sequences. In this case, the true positive rate (TPR) is the fraction of TCR sequences that receive a distance score below a given threshold that bind the same epitope. The false positive rate (FPR) is the fraction of those that do not. From figure 4, it is clear that the TCR sequences from Dash dataset are easier to group based on their epitope than the TCR sequences of the VDJdb dataset. As the TPR and FPR are independent from the size of the dataset and the negative/positive fraction, this is not due to the larger size of the VDJdb dataset. The reason for this different performance may be due to the fact that the VDJdb dataset is derived from a more diverse group of individuals or the assignments may be of a lower quality as only the most unreliable assignments were excluded. For the Dash dataset, it is clear that all methods are able to provide better distance scores to TCR sequences that bind the same epitope than would be expected at random. This includes both the Length and VJedit distance measures. Thus, in this dataset, TCR sequences with similar CDR3 length or V/J assignments bind similar epitopes. Furthermore, the GapAlign, Levenshtein and Profile methods are able to achieve around 40% TPR without any false positives. The Trimer and Dimer methods also perform strongly and exceed the GapAlign and Profile methods at higher FPR values. The ROC curve seems to indicate that the Trimer and Dimer methods are less stringent for exact matches but may be more capable to group very distant TCR sequences that do still bind the same epitope. For the VDJdb dataset, all methods do perform better than random, except for the VJedit distance, which runs along the diagonal. However, the performance of each method does not seem very viable for actual usage. For any method at any threshold, we can expect a high number of false positives.

**Figure 4:**
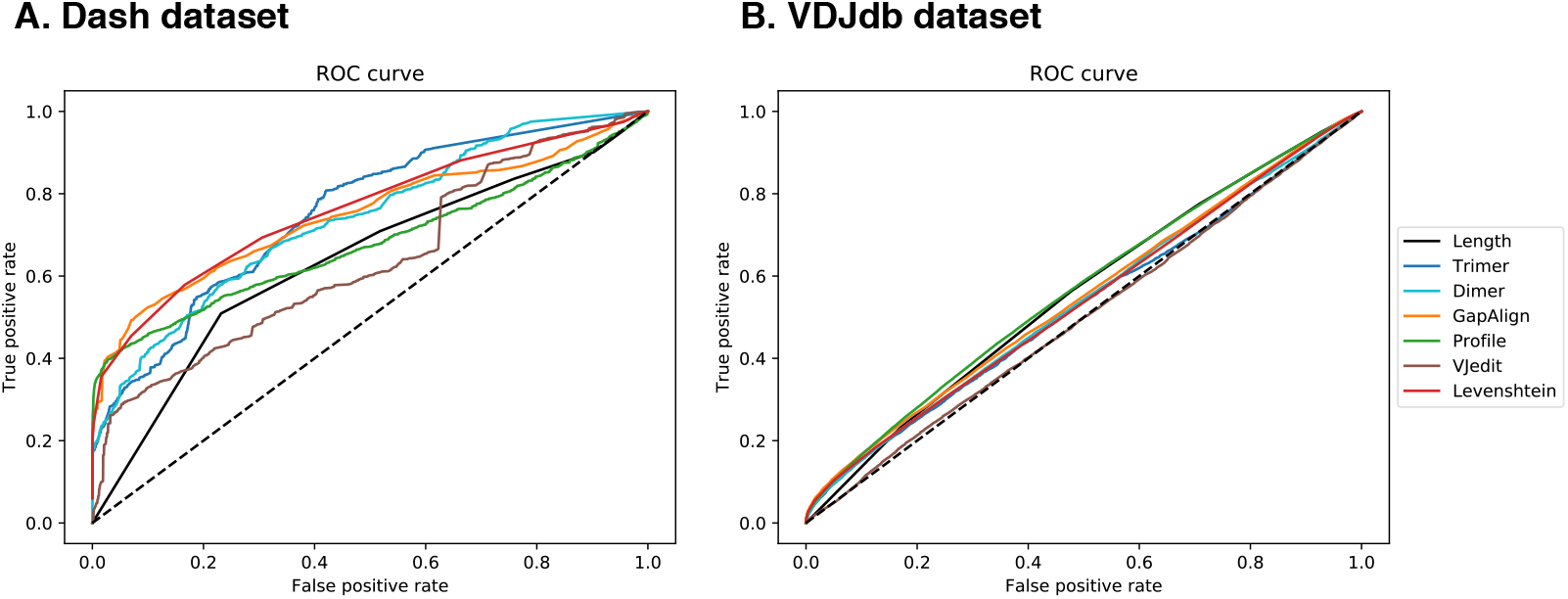
ROC plots for each of the six distance measures for the Dash dataset (A) and the VDJdb dataset (B). A low distance score for two TCR sequences that bind the same epitope is considered a true positive. A low distance score for two TCR sequences that do not bind the same epitope is then a false positive.

In a second analysis, we use the five distance measures as input for the DBSCAN clustering algorithm and evaluate the quality of the clusters at different thresholds. As can be seen in figure 5A, all methods have a strong performance on the Dash dataset. However, the random baseline is also high, since the majority (66.6%) of TCR sequences bind the same epitope, namely the M1 influenza epitope. Compared to this baseline, all methods feature a higher precision at most thresholds than random, except for the VJedit distance, which performs worse than random. Indeed, if half of the TCR sequences are grouped in a cluster (50% recall), all methods report that the clusters only consist of TCR sequences that bind the same epitope (100% precision). The accuracy, while high, is not as distinct from the random baseline for each method. This indicates that each method underperforms when having to group all TCR sequences that bind the same epitope into a single cluster. This suggests that each epitope has several groups of TCR sequences that are internally very similar and are easy to group with any distance measure. However, it seems much harder to bring together those different groups that target the same epitope.

**Figure 5:**
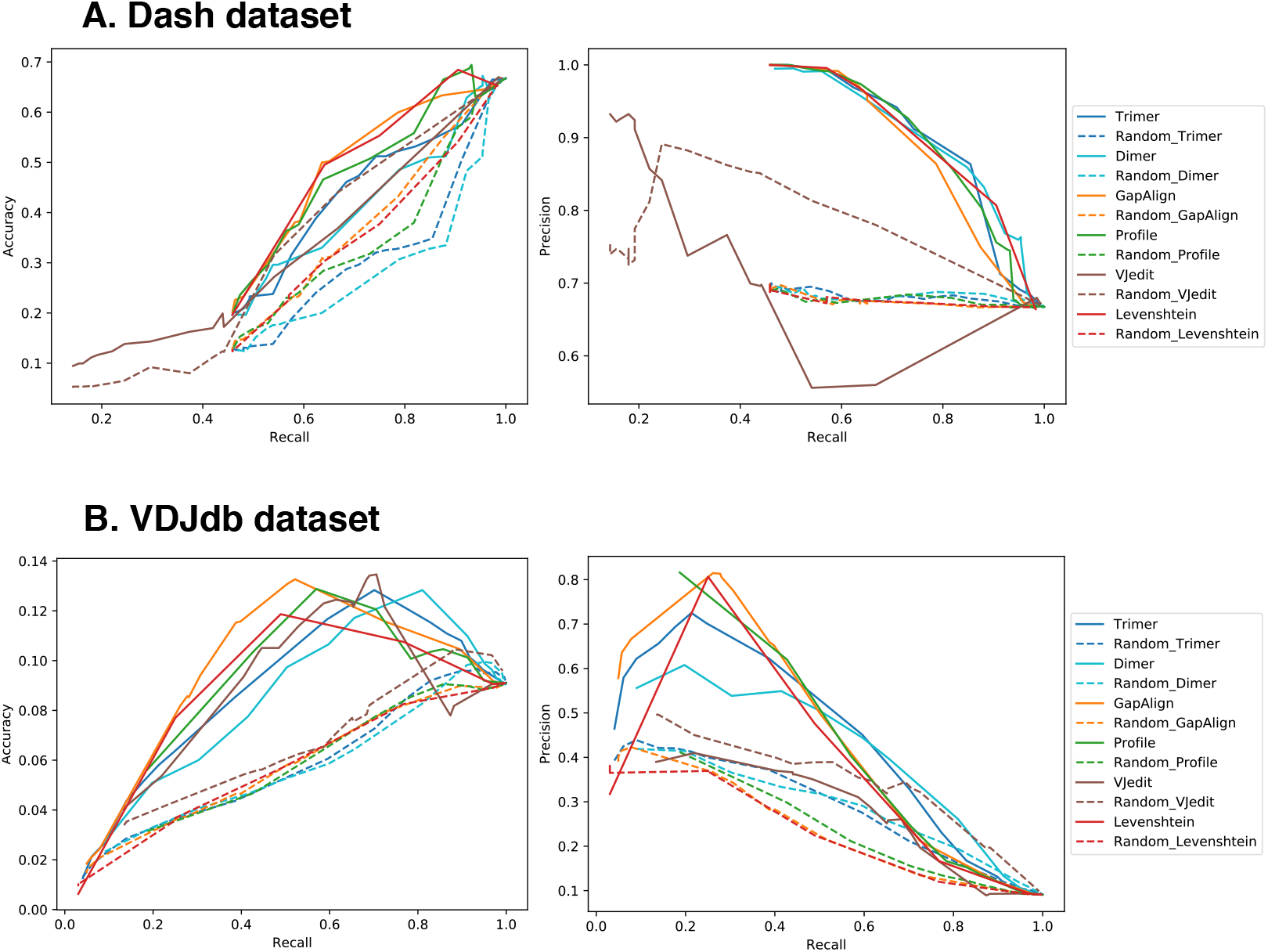
Accuracy-recall and precision-recall plots for each of the six distance measures for the Dash dataset (A) and the VDJdb dataset (B). The dotted lines denote the random baseline generated from 50 random permutations.

The VDJdb dataset reports similar trends as can be seen in figure 5B. In this case, there is a stronger distinction between the different methods. Furthermore, the precision at the highest recall drops down to 10% for all methods. The VJedit distance again performs worse than random. The GapAlign method performs best at the most stringent thresholds, where few TCR sequences are clustered but those that are clustered typically all bind the same epitope. At 80% precision, we can expect on average that for each cluster there will be one out of five TCR sequences that do not bind the same epitope as the others. The Levenshtein and Profile methods follow very similar trends to the GapAlign method. However, the Profile method is worse at the most stringent clustering cut-offs but achieves its highest performance at a slightly higher recall. The Trimer and Dimer methods follow the previously reported trend from the ROC curves, where they achieve their best performance when the cut-offs are less stringent. There is thus a clear decreasing stringency progression starting from GapAlign/Levenshtein, to Profile, to Trimer, to the Dimer method finally. In this manner, GapAlign is better for grouping the most similar TCR sequences together with high precision, Dimer is better at grouping very dissimilar TCR sequences with poor precision.

### Clustering thresholds are transferable between datasets

In the previous section, we screened the datasets for a large number of different clustering thresholds. However, in most practical applications, only one threshold will be used. As we have two datasets, we can use one to determine an optimal threshold and apply it to the other. The Dash dataset displayed high precision for all methods with a high recall. Thus, we selected the highest threshold for each method with the best precision for the Dash dataset and applied this threshold to the VDJdb dataset. The results can be found in table 1. For comparative purposes, we included the Length measure with a distance less than 1 so that all TCR sequences would be grouped by CDR3 length. As expected, the Length distance measure has a high recall as it can cluster all the TCR sequences without outliers, but has a poor accuracy and precision. On the other hand, the GapAlign, Profile, and Trimer measures all achieve high precision while clustering around a third of the dataset. This signifies that thresholds for these distance measures can be easily transposed between datasets, even if they have different sizes and densities. In addition, the most optimal threshold for the Levenshtein measure is a value of one, signifying a single amino acid change. This Levenshtein threshold value resulted in a quarter of the dataset being clustered, with the highest reported precision of all methods. Thus, the noted groups of similar TCR sequences that target the same epitope typically differ only by a single amino acid in their CDR3 sequence. The other methods are able to group a large subset of the available TCR sequences but only at the cost of lower precision, thus more TCR sequences that target different epitopes within a single group.

**Table 1:** Clustering statistics on the VDJdb with a fixed threshold

| Method | Threshold | Recall | Accuracy | Precision |
| --- | --- | --- | --- | --- |
| Length | 0.5 | 100% | 14.7% | 15.2% |
| Levenshtein | 1 | 24.8% | 7.7% | 80.6% |
| GapAlign | 18 | 30.7% | 9.4% | 77.6% |
| Profile | 1 | 28.3% | 7.6% | 76.1% |
| Trimer | 0.34 | 34.1% | 7.9% | 65.4% |
| Dimer | 0.23 | 52.0% | 9.9% | 49.2% |
| VJedit | 0.408 | 69.4% | 10.2% | 19.0% |

### Similar longer TCR sequences are more likely to bind the same epitope

A precision of 75% indicates that on average each cluster contains three out of four TCR sequences that bind the same epitope. However as this is an average, it is worthwhile investigating if this is uniform across all clusters or if there are some clusters with high precision and others with low precision. For the purposes of this analysis, we used the GapAlign method as this distance measure has been established as one of the most stringent. At the set threshold of 18, we divided the found TCR sequence clusters into a so-called ‘good’ group and a so-called ‘bad’ group. The ‘good’ group are those clusters where all the TCR sequences target the same epitope. An example of a good cluster can be found in table 2. The ‘bad’ group are those clusters that remain, thus where at least one TCR sequence recognizes other epitopes than the remainder. An example of a bad cluster can be found in table 3. Through this division, we found that indeed the majority of the clusters have a high precision and thus fall in the ‘good’ category. A minority of the clusters fall in the ‘bad’ category and have a very poor precision. The precision distribution is thus bimodal, as shown in figure 6. The question then becomes: “What makes a cluster ‘bad’?”. There is a clear set of TCR sequences that despite being highly similar, bind entirely different epitopes.

**Figure 6:**
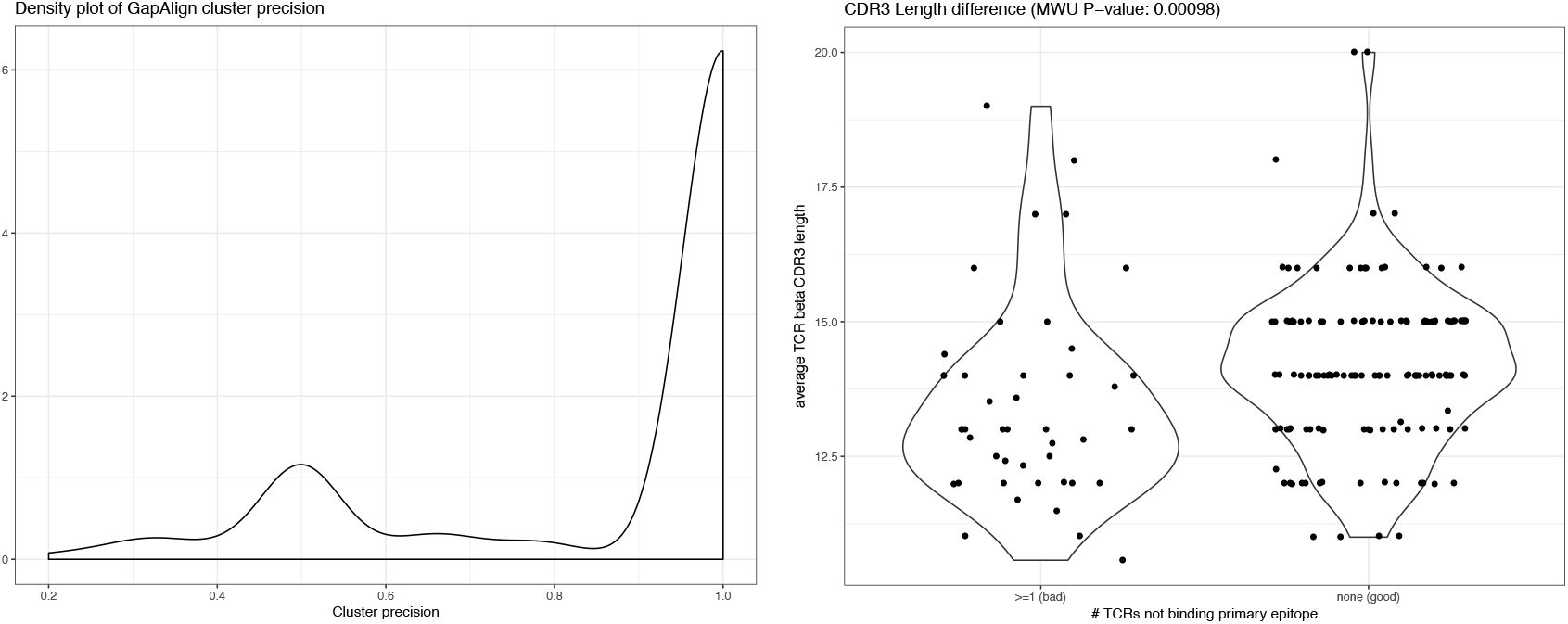
Left: Density plot of the precision for the GapAlign clusters found using DBSCAN at the fixed threshold of 18. Right: Violin plot of the TCRb CDR3 sequences lengths of the ‘bad’ clusters and the ‘good’ clusters. Each point represents a single cluster. A bad cluster is defined as containing at least one TCR sequence that binds a different epitope than all the others.

**Table 2:** Example of a ‘good’ cluster

| TCRb CDR3 Sequence | Epitope sequence |
| --- | --- |
| CASSEGRVSPGELFF | GLCTLVAML |
| CASSTGQVSPGELFF | GLCTLVAML |
| CASSEGQVSPGELFF | GLCTLVAML |
| CASSAGRVSPGELFF | GLCTLVAML |
| CASSEGRVLPGELFF | GLCTLVAML |
| CASSEGRISPGELFF | GLCTLVAML |
| CASSTGRVAPGELFF | GLCTLVAML |
| CASSEGRVLSGELFF | GLCTLVAML |

**Table 3:**
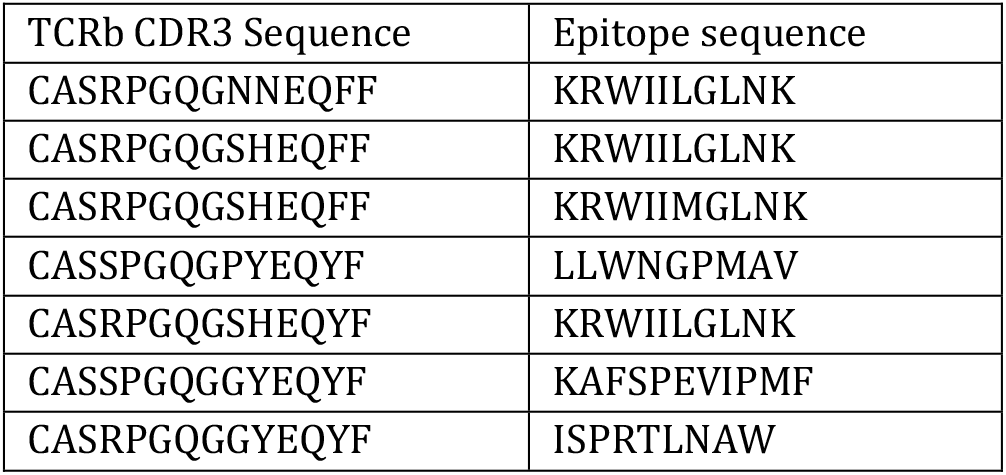
Example of a bad cluster

The properties of the CDR3 sequence of the TCR beta-chain within each cluster with both the good and bad groups were investigated. In this instance, we investigated the length of the CDR3 amino acid sequence, the percentage of hydrophobic residues and the percentage of charged residues. The physicochemical properties seemed especially relevant as it can be supposed that the CDR3 region needs to make contact with the epitope’s amino acids to facilitate the recognition, which can be aided by specific residues. However, we found no detectable difference in this frequency within the central CDR3 region (charged residue P-value: 1.00 & hydrophobic residue P-value: 0.85). A large difference was noted based on CDR3 amino acid length, as can be seen in figure 6. Shorter TCRb CDR3 sequences were more common in the ‘bad’ clusters. This indicates that if the sequence is short, there may not be sufficient information for the distance measures to separate those that bind different epitopes. In the same manner, clusters with long TCRb CDR3 sequences always concern a single epitope. Whether this is an artefact of the VDJdb dataset or represents a true biological division cannot be answered at this time.

### TCR distance does not represent epitope distance

T-cell receptor recognition is known to have a high degeneracy as one TCR can recognise a large number of similar epitope peptides. This may be problematic for the evaluation of our clusters as thus far we have only considered TCR sequences that bind exactly the same epitope. It may be the case that similar TCRs that are grouped bind very similar epitopes, but not the same one. Furthermore, the question can be raised that if similar TCRs bind similar epitopes, do dissimilar TCRs bind dissimilar epitopes. To this end, we plotted the distance between two TCR groups that bind two different epitopes against the distance between the epitopes themselves. For comparative purposes, we use the GapAlign method to calculate distances between the epitopes as it is closely related to traditional sequence alignment and the other methods cannot be readily transposed to the epitope space. As can be seen in figure 7, the overall correlation between the TCR sequence distance and the epitope distance is low. In fact in most cases, it is even negative. This seems to suggest that TCR sequences binding more similar epitopes have less similar TCR sequences.

**Figure 7:**
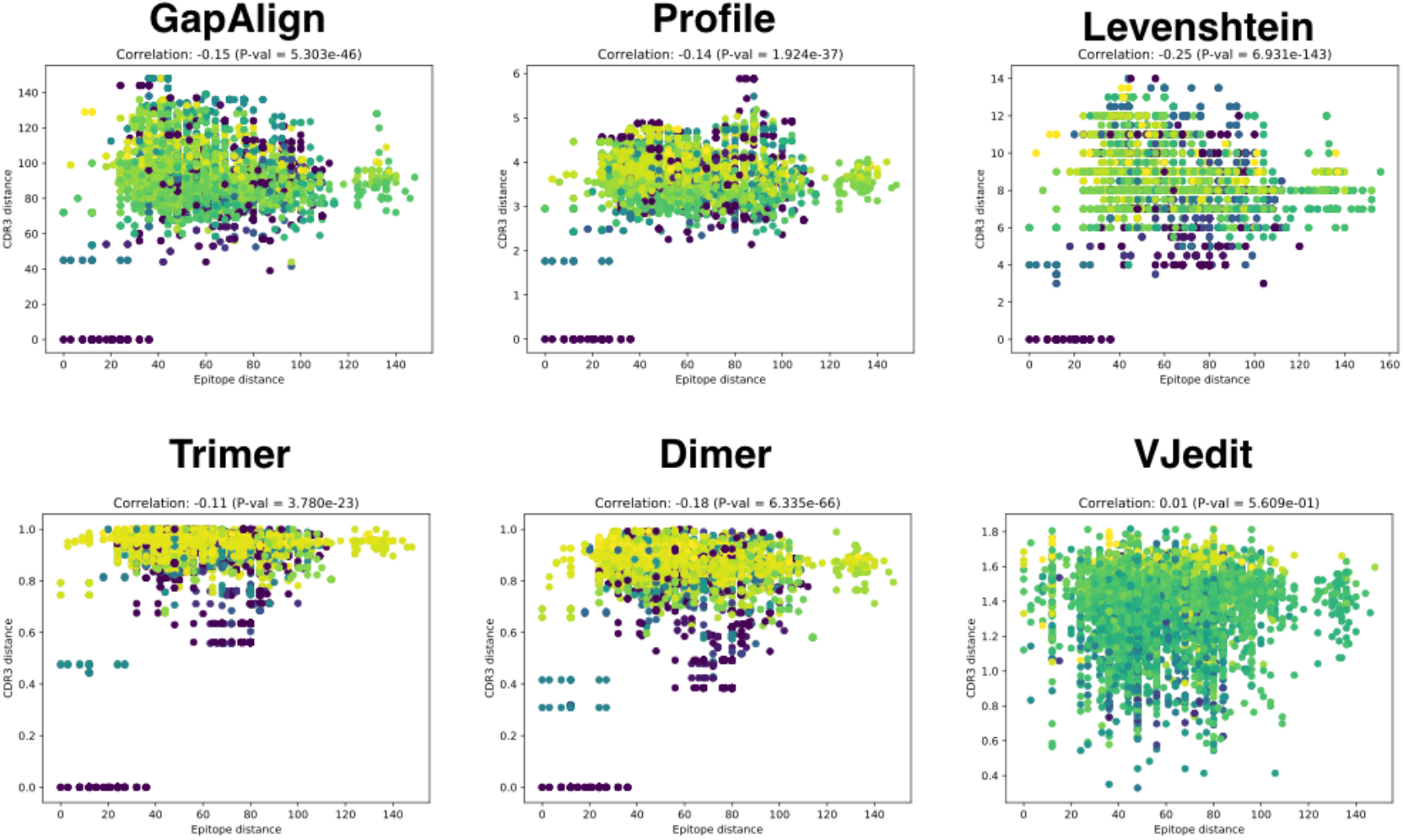
Scatter plots representing the relationship between TCR sequence distance and epitope sequence distance. Each point represents the comparison between a group of TCR sequences binding one epitope and a group of TCR sequences binding another epitope. On the y-axis the median distance between all TCR sequences from one group to the other is reported based on the specified distance measure. On the x-axis is the distance between the two epitopes. Reported are the Spearman correlations between the two measures. The colors indicate density at a given location in the plot, with the yellow color indicating higher density.

Furthermore, there are several epitope distances greater than a score of 120. These very dissimilar epitopes are those that originate from different MHC classes as they typically differ greatly in length (9.2 amino acids mean for MHC class I and 15.6 for MHC class II in the VDJ dataset). Thus, these represent comparisons between TCR sequences targeting a MHC class I epitope and TCR sequences targeting a MHC class II epitope. It is interesting to see that none of the distance measures provides large distances for TCR sequence groups that target different MHC classes. This signifies that using the definitions from these distance measures, there is no intrinsic difference in TCRs with different MHC class binding, despite the large epitope dissimilarity. This is unexpected as prior work has indicated significant differences within the CDR3 region between CD8+ and CD4+ T-cells (Li *et al*., 2016). Comparing the TCR CDR3 amino acid lengths assigned to either a peptide bound by MHC class I or MHC class I reveals no statistically significant difference (P-value= 0.49). This may be due to the much smaller size of this dataset compared to those used in the past.

However, for all distance measures, except for the VJedit, there are some comparisons with a median distance of 0, i.e. identical CDR3 sequences, and these typically target very similar epitopes. This may represent the degeneracy of the TCR, but they remain the exception rather than the rule. Detailed inspection reveals that the largest differences between TCR sequences are reported for those epitopes for which a large number of associated TCRs have been reported. In fact, the number of TCR sequences is a much better indication of the median distance than the epitope dissimilarity, as can be seen in figure 8. For example, the found Spearman correlation in the case of the Trimer distance measure is 0.27 (P-value = 1.99e-123) between the number of TCR sequences in the largest group and the overall median distance score. This again supports the presence of different dissimilar TCR groups that bind the same epitope. In addition, this suggests that epitopes with more known TCR sequences have a large diversity of such sequences. This may be due to data itself, where more sequences allow for large variation or these sequences are derived from distinct studies and populations, increasing variability. It may also have a biological reason, as epitopes that are more easily recognized by a large diversity of TCR molecules will have a much larger set of known TCR sequences revealed through tetramer staining followed by TCR sequencing.

**Figure 8:**
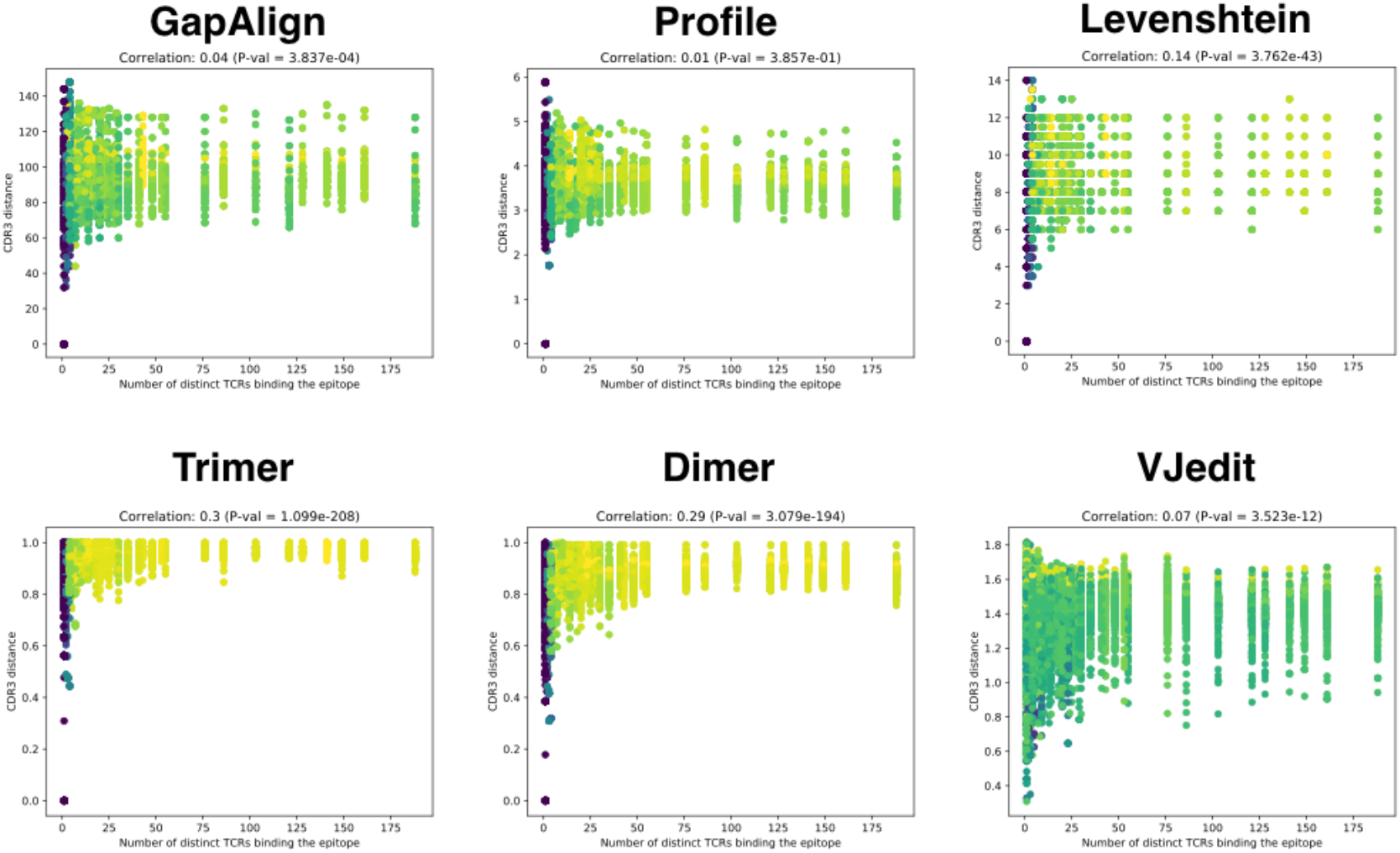
Scatter plots representing the relationship between TCR sequence distance and the number of TCR sequences associated with the epitope. Each point represents the comparison between a group of TCR sequences binding one epitope and a group of TCR sequences binding another epitope. On the y-axis the median distance between all TCR sequences from one group to the other is reported based on the specified distance measure. On the x-axis is the number of associated TCR sequences for the biggest group. Reported are the Spearman correlations between the two measures. The colors indicate density at a given location in the plot, with the yellow color indicating higher density.

## Conclusions

In this study, we investigated the performance of various unsupervised distance measures to group the beta-chain CDR3 amino acid sequence of TCR proteins with the aim to group those that recognize the same epitope. From our results, we can conclude that all distance measures using only the CDR3 region outperform random clustering, however none are able to perform perfect clustering of the entire dataset. Each measure had its own distinct advantages and disadvantages, but all performed comparably. A finding of interest in this manner was that a simple Levenshtein distance with a threshold of one was already highly performant and equivalent to many of the more advanced measures explored. Thus, one can achieve reasonable clustering of epitope-specific TCR sequences based on three simple criteria: 1) if they have identical length, 2) if the CDR3 amino sequence is sufficiently long and 3) if they differ by at most one amino acid. Clustering all TCR CDR3 amino acids targeting the same epitope is a much harder problem as they often end up in different distant clusters. Indeed, all distance measures agreed that there could be as much dissimilarity between two TCR CDR3 sequences targeting the same epitope, as two TCR CDR3 sequences targeting widely different epitopes presented by a different MHC class. Thus very different CDR3 beta-chain sequences can be associated with the same epitope, highly complicating any unsupervised or even a supervised approach. This suggests that despite the early successes with supervised methods, current techniques fall short of clustering all TCR sequences with the same unknown epitope preference in a robust and complete manner. Such techniques would however be useful to investigate naively sequenced repertoires, or T-cell collections where only the antigen protein is known. However, it should be noted that all conclusions in this study are highly dependent on the available dataset. There is no doubt that the future will see an expansion of TCR sequencing data along with novel experimental and computational techniques to process them. Finally, it should be noted that only the TCR beta sequence was used as input for the distance measures. Using both alpha- and beta-chain information may provide far superior performance, but there is still a substantial lack of this type of data within the scientific literature to evaluate this in a thorough manner.

